# UBA1 Mitigates Myocardial Ischemia/Reperfusion Injury by Attenuating Endoplasmic Reticulum-Mitochondria Contacts via Pdzd8 ubiquitination

**DOI:** 10.64898/2026.01.21.700963

**Authors:** Luo-Luo Xu, Pang-Bo Li, Wen-Xi Jiang, Jie Du, Hui-Hua Li

## Abstract

**BACKGROUND:** Myocardial ischemia/reperfusion injury (I/RI) represents a serious clinical complication in patients after acute myocardial infarction. Ubiquitin-activating enzyme 1 (UBA1) catalyzes the initial step of ubiquitination and plays a fundamental role in regulating protein homeostasis and related diseases. This study aims to elucidate the functional contribution of UBA1 to the pathogenesis of myocardial I/RI and to uncover its underlying mechanisms.

**METHODS:** Single-cell RNA sequencing was employed to characterize UBA1 expression in human ischemic heart tissues. Myocardial I/R injury was examined in myocardial-specific UBA1 knockout (UBA1^cko^) mice, UBA1-overexpressing mice (rAAV9-UBA1), and corresponding controls. Neonatal rat cardiomyocytes underwent hypoxia/reoxygenation *in vitro*. Cardiac function and infarction were evaluated by echocardiography and pathological staining. Protein–protein interactions were analyzed via immunoprecipitation combined with mass spectrometry. The endoplasmic reticulum–mitochondrial contact sites (ERMCSs) and mitochondrial ultrastructure were evaluated through transmission electron microscopy and confocal imaging.

**RESULTS:** UBA1 expression was significantly downregulated in human and murine ischemic myocardium, especially in cardiomyocytes. UBA1^cko^ mice exhibited aggravated I/RI with greater infarct size, impaired function, apoptosis, elevated intracellular Ca^2+^ levels, mitochondrial dysfunction, and ER stress, whereas UBA1 overexpression conferred cardioprotective effects. Mechanistically, UBA1 directly bound to and ubiquitinated Pdzd8, a key ERMCS-tethering protein, thereby promoting its degradation, which inhibited ERMCS formation and improved mitochondrial dysfunction and ER stress. Moreover, knockdown of Pdzd8 via rAAV9-siRNA effectively mitigated UBA1 knockout-induced myocardial damage. Additionally, administration of auranofin (AF), a U.S. Food and Drug Administration-approved drug for treating rheumatoid arthritis, markedly alleviated myocardial I/RI via activating UBA1 *in vivo* and *in vitro*.

**CONCLUSIONS:** UBA1 confers protection against myocardial I/RI by limiting ERMCS formation through Pdzd8 ubiquitination. Activating UBA1 or targeting Pdzd8 as a potential therapeutic strategy for the treatment of ischemic heart disease.

**GRAPHIC ABSTRACT:** A graphic abstract is available for this article.

**Clinical Perspective:** *What Is New?:*

- UBA1 expression is downregulated in human and murine ischemic myocardium, especially in cardiomyocytes.
- Cardiac deletion of UBA1 significantly exacerbates myocardial ischemia/reperfusion injury (I/RI), whereas cardiac UBA1 overexpression confers a marked protective effect.
- UBA1 interacts with Pdzd8 (PDZ domain containing 8) and facilitates its ubiquitination and subsequent degradation, which then reduces endoplasmic reticulum-mitochondria contact sites (ERMCSs) and ameliorates mitochondrial dysfunction and ER stress, protecting myocardial I/RI.
- Pharmacological activation of UBA1 with the FDA-approved drug auranofin attenuates myocardial I/R injury and improves heart dysfunction.

*What Are the Clinical Implications?:*

- UBA1 represents a new therapeutic target for myocardial I/RI.
- Activating UBA1 or targeting Pdzd8 may offer a promising therapeutic strategy for mitigating myocardial I/RI and heart failure, underscoring its potential for clinical translation.

## INTRODUCTION

Ischemic heart disease is a major cause of cardiac morbidity and mortality worldwide.^1^ Myocardial ischemia–reperfusion injury (I/RI) remains a major complication in patients undergoing angioplasty or thrombolytic therapy for acute myocardial infarction (MI).^2^ Multiple pathological processes, including inflammation, reactive oxygen species (ROS) accumulation, calcium overload, mitochondrial dysfunction, and endoplasmic reticulum (ER) stress, contribute to cardiomyocyte death, including apoptosis, necrosis, ferroptosis, necroptosis, and pyroptosis, during I/RI.^3,4^ Therefore, understanding the molecular mechanisms driving these events is essential for identifying new therapeutic targets.

Post-translational modification is a key regulator of protein function and signaling in various types of cells. Ubiquitin (Ub) and ubiquitin-like proteins (UBLs) control diverse cellular processes by altering protein stability, activity, localization, and trafficking. Ub/UBL conjugation is initiated by adenosine triphosphate (ATP)-dependent E1 activating enzymes and refined through a multi-step enzymatic cascade.^5^ Eight E1s have been identified, including those mediating ubiquitylation (UBA1, UBA6), SUMOylation (SAE1/SAE2), neddylation (NAE1), URMylation (UBA4), UFMylation (UBA5), and autophagy (ATG7).^6^ Ubiquitination eliminates misfolded or damaged proteins through coordinated actions of E1s, E2s, E3 ligases, and deubiquitinases (DUBs).^7^ E1s sit at the apex of this cascade, catalyzing ubiquitin activation to tag proteins for degradation.^6^ Of the two human E1s, UBA1 (UBE1) is essential for development, cell growth, and protein homeostasis. Its deletion is embryonic lethal in mice,^8^ and somatic UBA1 mutations cause VEXAS syndrome, a severe adult-onset autoinflammatory disorder.^9^ UBA1 also regulates apoptosis, neurodegeneration, and tumor biology by targeting diverse substrates.^10,11^ Dysregulated UBA1 expression has been linked to some cardiovascular disorders, including atherosclerosis, cardiac remodeling, and restenosis,^12–14^ yet its role in myocardial I/RI remains unclear.

Mitochondria and the ER are central regulators of cellular survival and homeostasis. Emerging studies show that mitochondria-associated ER membranes (MAMs) or ER–mitochondria contact sites (ERMCSs) act as communication hubs that regulate calcium transfer, lipid exchange, ROS homeostasis, mitochondrial dynamics, and ER stress.^15,16^ Several tethering proteins maintain ERMCS structure and function, including PDZ domain–containing 8 (Pdzd8), inositol 1,4,5-triphosphate receptor (IP3R), voltage-dependent anion channel 1 (VDAC1), 75-kDa glucose-regulated protein (GRP75), protein tyrosine phosphatase–interacting protein-51 (PTPIP51), diaphanous-1 (DIAPH1), and Mfn2. Dysregulation of these proteins leads to ERMCS abnormalities that contribute to diseases, such as aging-related disorders, neurodegeneration, metabolic syndrome, cancer, and cardiac pathology.^17–20^ However, the molecular mechanism regulating these tethering proteins and ERMCS function during myocardial I/RI remains largely unknown.

In this study, we found reduced UBA1 expression in hearts from MI patients, I/R mice, and hypoxia/reoxygenation (H/R)-treated cardiomyocytes. Cardiac-specific UBA1 knockdown worsened, whereas UBA1 overexpression improved I/R-induced myocardial injury and dysfunction. We also identified UBA1 binding to Pdzd8 and promoting its ubiquitination and proteasomal degradation, thereby reducing ERMCSs formation and limiting mitochondrial dysfunction and ER stress. Importantly, administration of the UBA1 activator auranofin further prevented I/R-induced increases in ERMCSs and myocardial injury. These findings identify the UBA1-Pdzd8 axis as a mechanism contributing to myocardial I/RI and highlight this pathway as a potential therapeutic target.

## METHODS

Detailed experimental methods and Major Resources Table are available in the Supplementary Material. The data for the figures are not shown and are available from the corresponding author upon reasonable request. RNA sequencing data have been deposited in the National Center for Biotechnology Information Gene Expression Omnibus (GSE145154).

The role of UBA1 in myocardial infarct size (I/RI) was evaluated in UBA1^CKO^ mice and rAAV9-UBA1 mice, respectively, subjected to 30 min of ligation of the left anterior descending coronary artery, followed by a 24-h release of the ligature. Echocardiography, histological analysis, and transmission electron microscopy were performed to assess the function of UBA1 in myocardial function and injury after I/R surgery. All animal protocols in this study were approved by the Institutional Animal Care and Use Committee of Chaoyang Hospital (2020-Animal-164) of Capital Medical University and were performed following the criteria of the Guide for the Care and complied with the National Institutes of Health Guide for the Care and Use of Laboratory Animals (1996).

The effects of UBA1 on cardiomyocyte ERMCSs formation and hypoxia-driven injury were investigated *in vitro* using neotatal rat cardiomyocytes (NRCMs) with adenovirus-mediated UBA1 silencing or overexpression. RNA sequencing, immunoprecipitation coupled with mass spectrometry, coimmunoprecipitation, protein pull-down, confocal imaging, ubiquitination assays, and Oxygraph-2k analysis were performed to clarify the molecular mechanisms of UBA1 in ERMCSs formation and myocardial I/RI.

### Statistical Analysis

All data are presented as mean ± standard error of the mean (SEM). Statistical analyses were performed with GraphPad Prism v10.3.1. Normality was assessed by the Shapiro–Wilk test. Comparisons between two groups were carried out with the paired or unpaired Student’s t-test or the appropriate non-parametric counterpart, chosen according to data distribution. Multiple-group comparisons were evaluated by one-way or two-way ANOVA followed by Tukey’s post-hoc test. Longitudinal group differences were analysed with linear mixed-effects models. Associations between variables were examined using Pearson’s correlation. Results are reported as mean ± SEM. A P value less than 0.05 was considered significant.

## RESULTS

### Upregulation of UBA1 in Mouse I/R Hearts and H/R Cardiomyocytes, and MI Patient Hearts

To evaluate whether Ub/UBL-mediated protein modification contributes to myocardial I/RI, we analyzed differentially expressed genes (DEGs), focusing on E1 enzymes, in human normal and ischemic heart samples using a published single-cell RNA sequencing (scRNA-seq) dataset (GSE145154) (Figure 1A). Four E1 candidates (UBA1, UBA5, UBA6, and ATG7) were altered in cardiomyocytes, with UBA1 showing the most pronounced downregulation in ischemic hearts compared with controls (Figure 1B). We next confirmed this in a mouse I/R model (Figure 1C). qPCR showed time-dependent reductions in UBA1 mRNA in I/R hearts of wild-type (WT) mice (Figure 1D), and UBA1 protein downregulation was validated by immunoblotting and immunohistochemistry (Figure 1E and 1F). We then used an in vitro H/R model in neonatal rat cardiomyocytes (NRCMs) and fibroblasts (NRCFs) (Figure 1G). UBA1 protein levels declined over time in NRCMs, but not NRCFs, after 24 h H/R stimulation (Figure 1H), supporting cardiomyocyte-specific reduction.

**Figure 1.**
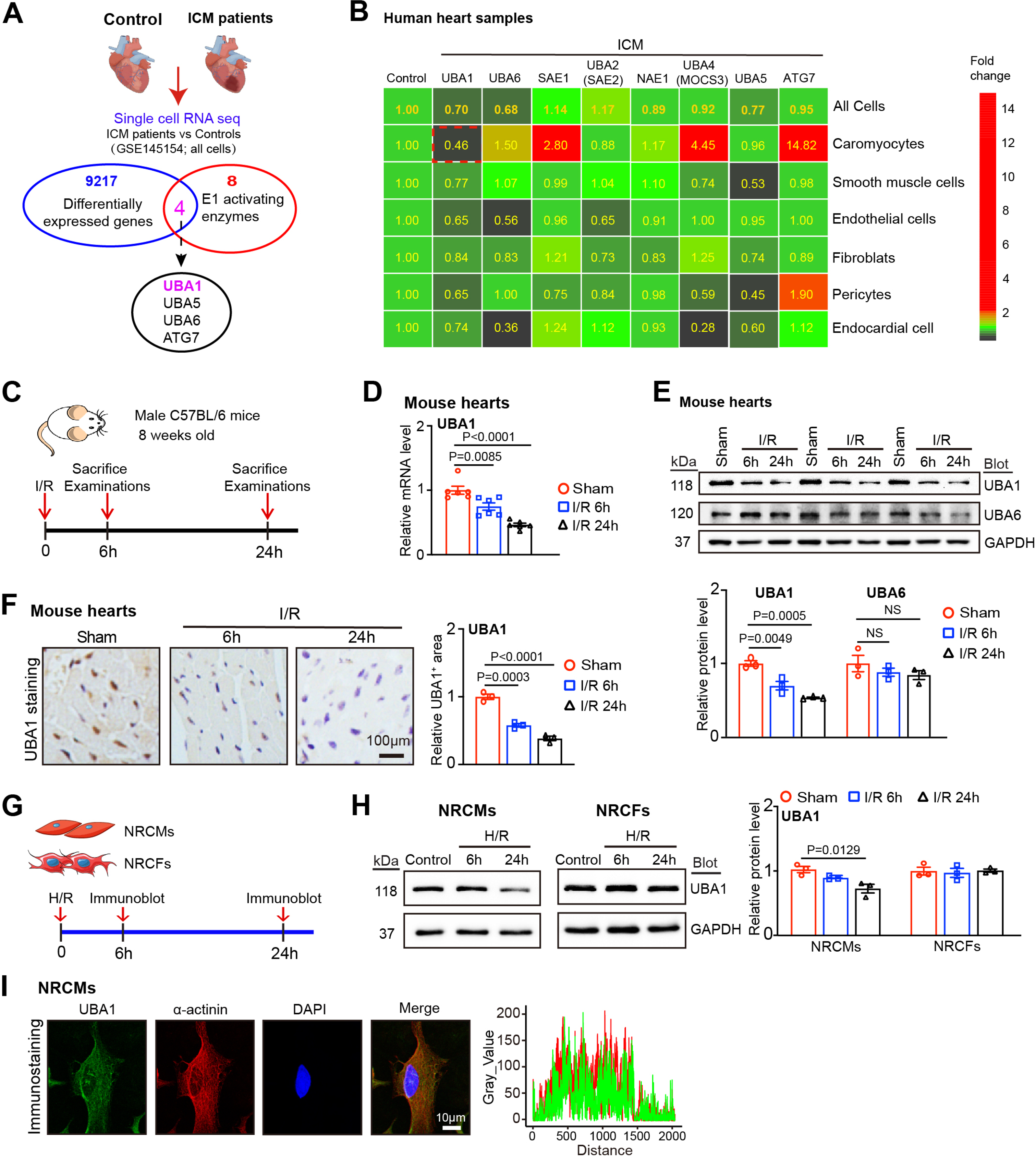
Reduction of UBA1 expression in mice with ischemia/reperfusion (I/R) and patients with myocardial infarction. **A,** Venn diagram showing the overlap of differentially expressed genes identified from a single-cell RNA sequencing (sc-RNA seq) dataset (GSE145154) of ICM patients and healthy controls, leading to the identification of four key E1 activating enzymes. **B,** The heatmap of the expression levels of ubiquitin-like modifier activating enzyme-related genes (UBA1, UBA5, UBA6, SAE1, SAE2, NAE1, MOCS3 and ATG7) from human ischemic hearts examined by sc-RNA seq dataset (GSE145154). **C,** Schematic of the I/R experiment in 8-week-old male C57BL/6 mice. Examinations were performed at 6 h and 24 h after reperfusion. **D,** qPCR analysis of the UBA1 mRNA levels in the broad zone of the heart (n = 6). **E,** Immunoblotting analysis of UBA1 protein level in the border zone of the heart (upper) and quantification of the protein levels (lower, n = 4). **F,** Immunohistochemical staining of heart slices with an anti-UBA1 antibody (left) and quantification of the UBA1 area (right, n = 6). Bar: 100 μm. **G,** Neonatal rat cardiomyocytes (NRCMs) and neonatal rat cardiac fibroblasts (NRCFs) were exposed to hypoxia/reoxygenation (H/R) for 6 and 24 h, respectively. **H,** Immunoblotting analysis of UBA1 protein levels in H/R-treated NRCMs (left) and the quantification (right, n = 4). **I,** Fluorescence localization of the UBA1 and α-actinin proteins in NRCMs. Bar: 10 μm. Data are expressed as the mean ± SEM, and n indicates the sample number per group. Statistical significance was assessed by One-way ANOVA. *P* values are indicated in the figure.

Immunostaining further showed UBA1 localization at the Z-disc in NRCMs, marked by ɑ-actinin (Figure 1I). Collectively, these data indicate that UBA1 expression decreases in ischemic cardiomyocytes and may contribute to myocardial I/RI.

### Myocardial-specific depletion of UBA1 aggravates I/R-induced cardiac dysfunction in mice

To define the role of UBA1 in myocardial I/RI, we generated cardiac-specific UBA1 knockout (UBA1^cko^) mice by crossing UBA1^f/f^ mice with α-MHC–Cre mice. UBA1^f/f^ mice were healthy and used as controls (Figure S1A through S1D). Eight-week-old UBA1^f/f^ mice were administered tamoxifen (40 mg/kg) to induce UBA1 deletion. Five days (Figure S1E) later, UBA1 protein levels were reduced by ∼45% in UBA1^cko^ hearts but remained unchanged in other organs. Baseline cardiac structure and function were similar between groups. Mice then underwent sham or I/R surgery for 24 h (Figure 2A). UBA1 protein remained lower in UBA1^cko^ hearts and declined further after I/RI (Figure 2B). Echocardiography showed worse myocardial function in UBA1^cko^ mice, with greater reductions in ejection fraction (EF%) and fractional shortening (FS%) compared with UBA1^f/f^ controls (Figure 2C). I/R also increased LV end-systolic diameter (LVESD) and end-systolic volume (LVESV) in UBA1^f/f^ mice, with further enlargement in UBA1^cko^ mice (Table S3). UBA1^cko^ mice displayed larger infarcts, more TUNEL-positive cardiomyocytes, and higher Bax/Bcl-2 ratios (Figure 2D through 2F). Bax and serum lactate dehydrogenase (LDH) were increased, and Bcl-2 was reduced in UBA1^cko^ hearts after I/R (Figure 2G and 2H). ROS production (DHE staining) and NOX2/NOX4 expression were also elevated in UBA1^cko^ hearts (Figure 2I and 2J). Under sham conditions, function and pathology were comparable between groups. These findings suggest that UBA1 depletion accelerates I/R-induced myocardial injury and dysfunction.

**Figure 2.**
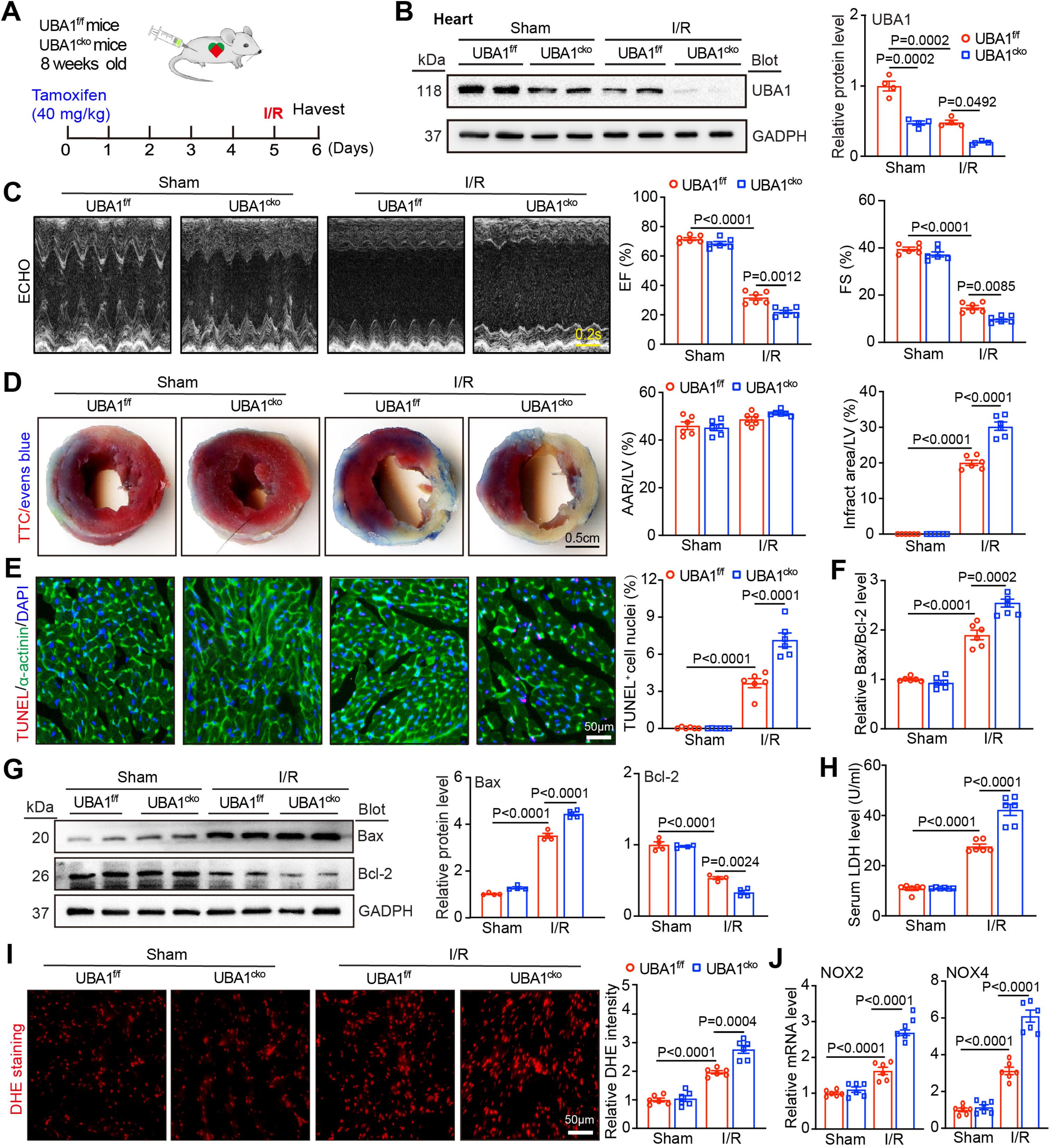
Cardiac-specific knockdown of UBA1 accelerates I/R-mediated cardiac injury and dysfunction. **A,** Schematic diagram of the experimental design for intraperitoneal injection of Tamoxifen (TAM; 40 mg/kg) and I/R surgery in male UBA1^f/f^ and UBA1^cko^ mice. **B,** Immunoblotting analysis of UBA1 knockdown efficiency in UBA1^f/f^ and UBA1^cko^ mice (n = 4). **C,** Echocardiographic (ECHO) examination of the left ventricle (LV) (left) and percentages of the ejection fraction (EF%) and fraction shortening (FS%) (right, n = 6). Bar: 0.2s. **D,** Representative images of heart sections stained with TTC-Evans blue dye (left). Percentages of the area at risk (AAR) to the LV area or the infarct area to the LV area (right, n = 6). Bar: 0.5 cm. **E,** Representative images of heart sections stained with TUNEL (red), anti-α-actinin (green) and DAPI (blue; left) and the percentages of TUNEL-positive nuclei (right, n = 6). Bar: 50 μm. **F,** qPCR analysis of Bcl-2 and Bax mRNA levels (left) and quantification of Bcl-2 to Bax ratio (right, n = 6). **G,** Immunoblotting analysis of Bcl-2 and Bax protein levels (left) and quantification (right, n = 4). **H,** Measurement of serum LDH activity in the border zone of ischaemic heart tissues by LDH assay kit (n = 6). **I,** Fluorescence staining heart sections with dihydroethidium (DHE) dye (left), and quantification of ROS levels (right, n = 6). Bar: 50 μm. **J,** qPCR analysis of NOX2 and NOX4 mRNA levels (n = 6). Data are expressed as the mean ± SEM, and n indicates the sample number per group. Statistical significance was assessed by two-way ANOVA. *P* values are indicated in the figure.

### Cardiac-specific knockdown of UBA1 enhances mitochondrial abnormalities and ER stress in mice

To investigate pathways mediating the increased susceptibility of UBA1^cko^ mice to I/R injury, we performed RNA-seq on hearts from UBA1^cko^ and UBA1^f/f^ mice 24 h after sham or I/R surgery. UBA1^f/f^ hearts showed 169 upregulated and 380 downregulated genes after I/R compared with sham, whereas UBA1^cko^ hearts showed 2419 upregulated and 2092 downregulated genes relative to UBA1^f/f^ controls (Figure 3A through 3C). Gene ontology (GO) and Wiki pathway analyses revealed marked reductions in pathways associated with mitochondrial protein complexes, the electron-transport chain (ETC), oxidative phosphorylation, and the tricarboxylic acid (TCA) cycle in UBA1^cko^ hearts after I/R (Figure 3D). Gene set enrichment analysis (GSEA) similarly identified the TCA pathway as strongly linked to UBA1 deficiency (Figure 3E). Heatmap analysis showed that I/R-induced decreases in genes related to mitochondrial complexes and TCA metabolism in UBA1^f/f^ hearts were further exacerbated in UBA1^cko^ hearts (Figure 3F). These findings indicate that loss of UBA1 markedly impairs mitochondrial metabolic programs during I/R.

**Figure 3.**
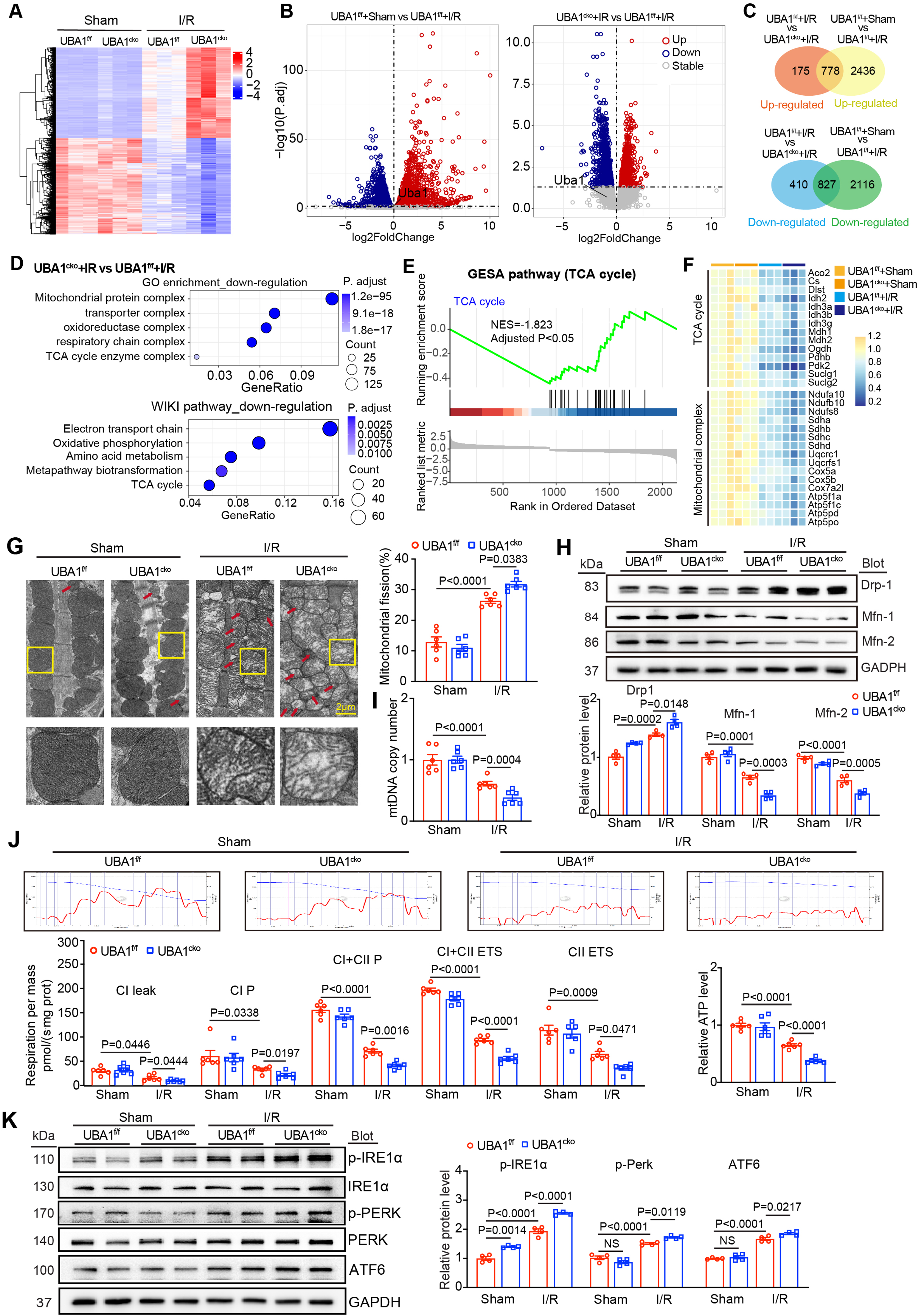
Cardiac-specific knockdown of UBA1 improves I/R-induced myocardial mitochondrial dysfunction and ER stress. **A,** RNA sequencing analysis of heart tissues and the heatmap of differentially expressed genes (DEGs) between UBA1^f/f^ and UBA1^cko^ mice after sham or I/R (n = 3). Red indicates upregulation; blue indicates downregulation. **B,** Volcano plot of DEGs between UBA1^f/f^ and UBA1^cko^ mice after sham or I/R. **C,** Venn diagram of DEGs between UBA1^f/f^ and UBA1^cko^ mice after sham or I/R. **D,** GO enrichment and WikiPathway analysis of the downregulated genes in UBA1^f/f^ and UBA1^cko^ mice post-I/R. **E,** Gene enrichment analysis (GESA) diagram of DEPs in UBA1^cko^ mice after I/R. **F,** Heatmap of DEGs in UBA1^cko^ mice after I/R. **G,** Representative images of transmission electron microscopy (TEM) showing mitochondrial morphology in cardiomyocytes (red arrows indicate fragmented mitochondria) and quantitative analysis of fragmented mitochondria (n = 6). Bar: 1 cm. **H,** Immunoblotting analysis of Drp1, Mfn1, and Mfn2 protein levels in the heart (left), quantitative data of each protein (n = 4). **I,** Mitochondrial DNA copy numbers determined by qPCR analysis. β-globin level as control (n = 6). **J,** Heart mitochondrial function was measured by an Oroboros Oxygraph-2K (O2K). Representative traces of O_2_ concentration (blue line) and O_2_ flux normalized to tissue mass (red line) (top). Summary data from Oxygraph-2k measurements of respiratory capacity, including CI leak (complex I proton leak), CI P (complex I oxidative phosphorylation), CI + CII P, CI + CII ETS (electron transport system capacity), CII ETS (bottom left), and ATP production (bottom right, n = 6). **K,** Immunoblotting analysis of endoplasmic reticulum (ER) stress relative protein levels (phosphorylated (p)-IREɑ, IREɑ, p-PERK, PERK and ATF6) (left) and quantification (right, n = 4). Data are expressed as the mean ± SEM, and n indicates the sample number per group. Statistical significance was assessed by two-way ANOVA. *P* values are indicated in the figure.

To confirm these findings, we examined mitochondrial structure and function in the hearts. Transmission electron microscopy (TEM) showed increased mitochondrial fragmentation in UBA1^f/f^ mice after 24 h of I/RI, with further fragmentation in UBA1^cko^ hearts (Figure 3G). UBA1^cko^ hearts also showed greater ultrastructural defects, including reduced cristae density and disorganized cristae membranes (Figure 3G). Because mitochondrial morphology depends on fission and fusion mediators such as Drp1, MFF, Mid49, Mid51, and Mfn1/2,^21^ we examined their expression. I/R increased Drp1, MFF, Mid49, and Mid51 and reduced Mfn1 and Mfn2 in UBA1^f/f^ hearts, with these changes more pronounced in UBA1^cko^ hearts (Figure 3H and Figure S2A). Consistent with enhanced fission, mitochondrial DNA (mtDNA) copy number was lower in UBA1^cko^ hearts (Figure 3I). Because the balance of mitochondrial dynamics supports ETC function and ATP production,^22,23^ we measured oxygen consumption. UBA1^cko^ hearts showed reduced mitochondrial oxidative phosphorylation (OXPHOS), diminished complex I and II activities, decreased mitochondrial membrane potential (MMP), and lower ATP after I/R (Figure 3J). However, no significant alterations in all parameters for mitochondrial structure and function were observed in the sham groups (Figure 3G through 3J), indicating that UBA1 deletion causes mitochondrial dysfunction specifically during I/RI.

Meanwhile, GO and KEGG analyses showed that upregulated genes in UBA1^cko^ hearts after I/R were enriched in immune effector pathways, including leukocyte migration, activation, chemokine signaling, and ER stress–related pathways such as ERAD, protein folding, and ER protein processing (Figure S2B). mRNA levels of multiple cytokines and chemokines (Cxcl5, Cxcl10, Ccl2, Ccl4, Ccl7, and Cxcl12) and their receptors (Ccr1, Ccr2, and Ccr5) were markedly higher in UBA1^cko^ hearts (Figure S2C). We next examined ER homeostasis. Immunoblotting showed that I/R-induced increases in p-IREɑ, p-PERK, and ATF6 in UBA1^f/f^ hearts were further elevated in UBA1^cko^ hearts (Figure 3H), indicating enhanced ER stress after UBA1 loss. Together, these findings show that UBA1^cko^ aggravates myocardial I/RI by promoting mitochondrial dysfunction and ER stress *in vivo*.

### Overexpression of UBA1 in Cardiomyocytes Alleviates Cardiac Injury, Mitochondrial Dysfunction, and Excessive ER Stress in Mice

To evaluate whether UBA1 overexpression protects the heart during I/RI, WT mice received rAAV9-NC or rAAV9-UBA1 via tail vein injection (Figure 4A). Three weeks later, immunoblotting showed a 1.4-1.6-fold increase in UBA1 expression in rAAV9-UBA1 mice (Figure 4B). Compared with rAAV9-NC, rAAV9-UBA1 markedly improved cardiac function (higher EF% and FS%), reduced infarct size, decreased TUNEL^+^ myocytes, and lowered the Bax/Bcl-2 mRNA ratio after I/R (Figure 4C through 4F, Table S4). rAAV9-UBA1 mice also exhibited higher Bcl-2, lower Bax, reduced ROS, decreased NOX2/4 mRNAs, and reduced LDH activity post-I/R (Figure S3A through S3C). Increases in fragmented mitochondria, upregulation of Drp1, MFF, Mid49, and Mid51, and reductions in Mfn-1 and Mfn-2 and mtDNA copy number were all reversed by rAAV9-UBA1 (Figure 4J through 4I, Figure S3D). rAAV9-UBA1 also restored OXPHOS, ETC complex I and II activities, MMP, and ATP content post-I/R (Figure 4J). Finally, I/R-induced increases in p-IREɑ, p-PERK, and ATF6 were reduced in rAAV9-UBA1 mice (Figure 4K). Together, these findings show that UBA1 overexpression attenuates myocardial I/RI by improving mitochondrial function and limiting ER stress.

**Figure 4.**
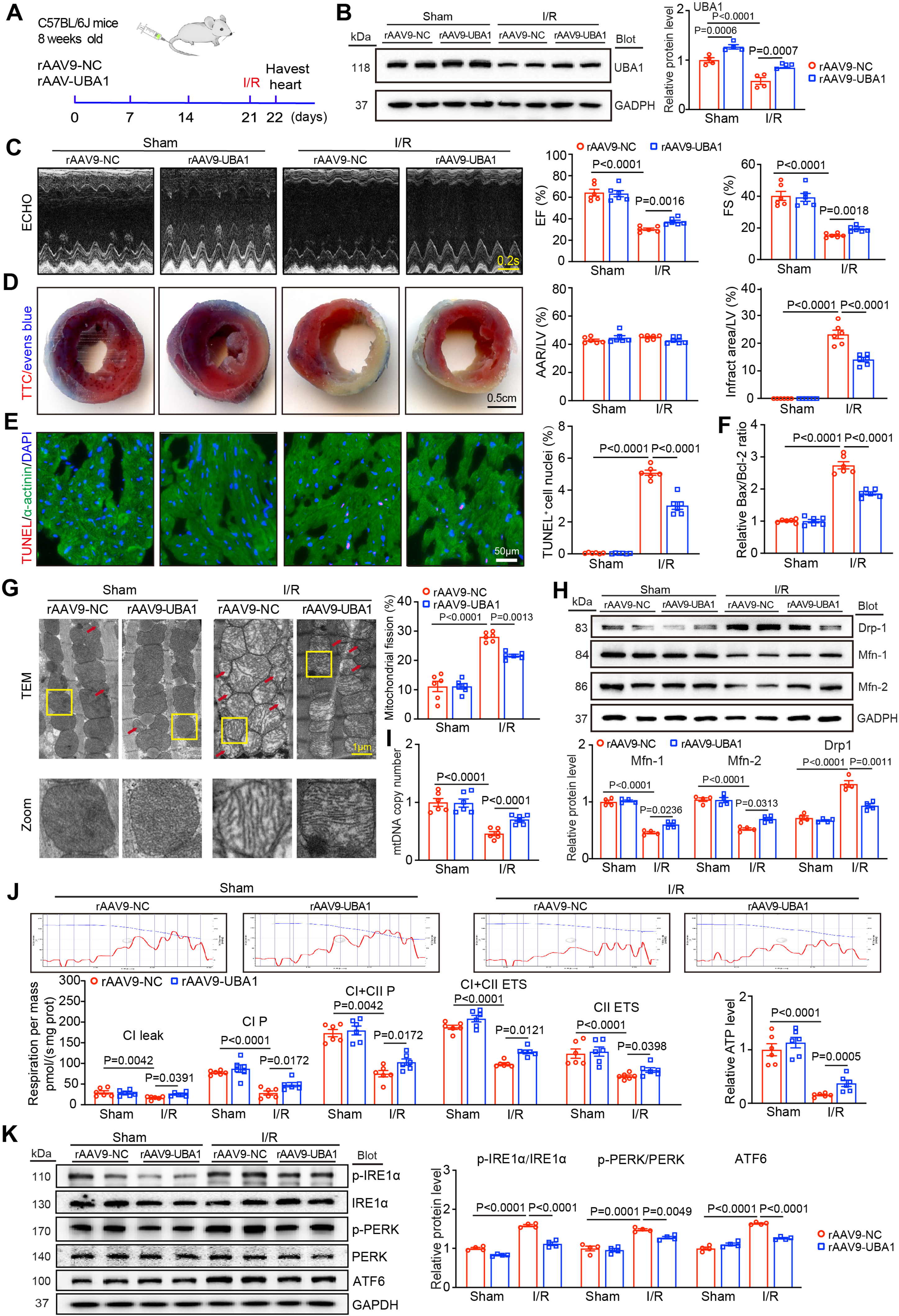
Cardiac-specific overexpression of UBA1 ameliorates I/R-induced myocardial mitochondrial dysfunction and ER stress. **A,** Schematic illustration of the UBA1-overexpressing mice injected by recombinant adeno-associated virus serotype 9 (rAAV9). **B,** Immunoblotting analysis of UBA1 overexpression efficiency in mice injected with rAAV9-UBA1 or rAAV9-NC control (n = 4). **C,** Representative echocardiographic images of left ventricular (LV) and quantification of EF% and FS% in rAAV9-NC- and rAAV9-UBA1-injected mice post-I/R (n = 6). Bar: 0.2 s. **D,** TTC–Evans blue staining of cardiac infarct areas, and the ratios of the AAR to the LV area and the infarct area relative to the LV area (n = 6). Bar: 0.5 cm. **E,** Representative images of TUNEL staining in border-zone of cardiac infarct areas (left). The percentage of TUNEL-positive nuclei (right, n = 6). Bar: 50 μm. **F,** qPCR analysis of Bax and Bcl-2 mRNA levels in the border zone of infarct areas (n = 6). **G,** Representative TEM images of cardiomyocyte mitochondrial morphology (red arrows: fragmented mitochondria) and quantification of fragmented mitochondria (n = 6). Bar: 1 cm. **H,** Immunoblotting analysis and quantification of Drp1, Mfn1 and Mfn2 protein levels in the heart (n = 4). **I,** qPCR analysis of ND-1 to determine mtDNA copy number, normalized to β-globin (n = 6). **J,** Representative traces of O_2_ concentration (blue line) and O_2_ flux normalized to tissue mass (red line) (top). Summary data from O2K measurements of respiratory capacity, including CI leak (complex I proton leak), CI P (complex I oxidative phosphorylation), CI + CII P, CI + CII ETS, CII ETS and ATP production (bottom, n = 6). **K,** Immunoblotting analysis of p-IREɑ, IREɑ, p-PERK, PERK and ATF6) (left) and quantification of p-IREɑ/IREɑ, p-PERK/PERK and ATF6 (right, n = 4). Data are expressed as the mean ± SEM, and n indicates the sample number per group. Statistical significance was assessed by two-way ANOVA. *P* values are indicated in the figure.

### UBA1 Regulates H/R-induced Cardiomyocyte apoptosis, Mitochondrial Dysfunction, and Activation of ER stress *in vitro*

To determine whether UBA1 knockdown worsens H/R-induced injury *in vitro,* NRCMs were transduced with control shRNA (Sh-NC) or shRNA targeting endogenous UBA1 (Sh-UBA1) and exposed to H/R for 24 h. Sh-UBA1 at MOI=50 reduced UBA1 by ∼50% without cytotoxicity (Figure S4A and 4B), so this dose was used. Consistent with *in vivo* results (Figure 2 and 3), UBA1 silencing increased TUNEL^+^ cells, ROS (DHE), Ca^2+^ release (Fluo4), and reduced MMP (Δψm, JC-1) after H/R (Figure S4C through 4F). UBA1 knockdown also enhanced mitochondrial fission, elevating Drp1, MFF, Mid49, and Mid51 and decreasing Mfn1 and Mfn2 (Figure S4G and 4H). ER stress markers p-IREα, p-PERK, and ATF6 were also higher in Sh-UBA1 cells (Figure S4I).

To confirm these findings, we tested whether UBA1 overexpression protects NRCMs. Ad-UBA1 at MOI = 50 increased UBA1 by ∼1.5-fold without cytotoxicity (Figure S5A and 5B). In contrast to Sh-UBA1, Ad-UBA1 reduced TUNEL^+^ cells, ROS, Ca²⁺, and mitochondrial fission, and increased Δψm after H/R (Figure S5C through 5G). UBA1 overexpression also restored mitochondrial dynamics and reduced ER stress, suppressing Drp1, p-IREα, p-PERK, and ATF6 while increasing Mfn1 and Mfn2 (Figure S5H). Under control conditions, neither UBA1 knockdown nor overexpression altered these parameters (Figure S5A through 5H). Overall, these results show that UBA1 overexpression protects cardiomyocytes from H/R injury by limiting mitochondrial dysfunction and excessive ER stress *in vitro*.

### UBA1 Interacts with Pdzd8 to Reduce Its Stability through Ubiquitination

To investigate how UBA1 regulates mitochondrial function and ER stress after I/R, we examined its interacting partners. Co-immunoprecipitation (Co-IP) with LC-MS/MS identified multiple proteins enriched by anti-UBA1 relative to immunoglobulin G (IgG) (Figure 5A through 5C, Table S5). Among these, Pdzd8, an ER transmembrane protein that regulates ERMCS formation, Ca^2+^ handling, mitochondrial homeostasis, and ER stress,^24,25^ was of particular interest (Figure S6A).

**Figure 5.**
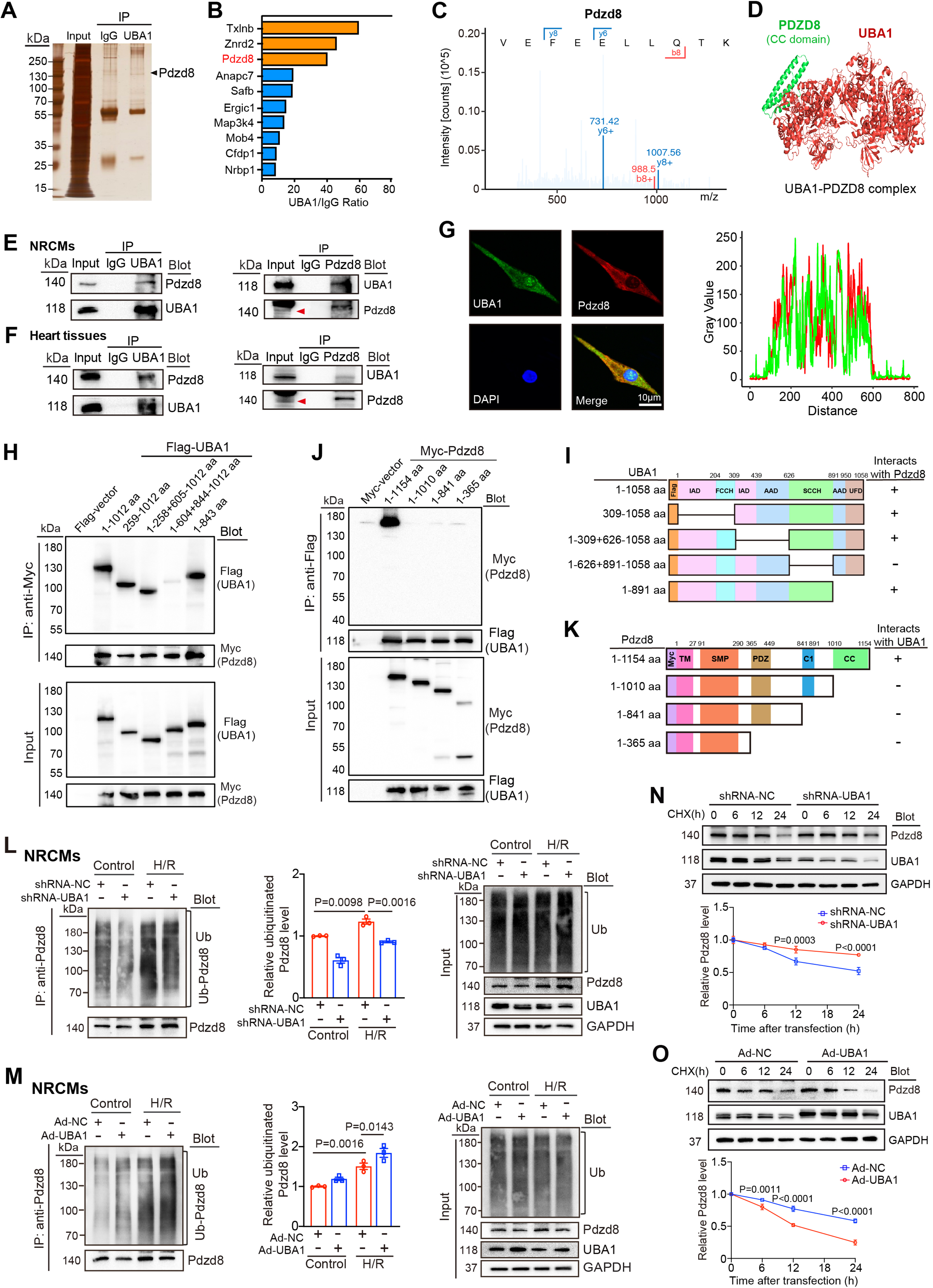
UBA1 directly interacts with Pdzd8 and promotes its ubiquitination. **A,** Silver-stained gel image of proteins obtained following UBA1 immunoprecipitation (IP) for protein-quality assessment. **B,** Top ten proteins ranked by interaction strength with UBA1 as determined by LC-MS/MS analysis after UBA1 IP. **C,** Representative MS/MS spectrum and extracted-ion chromatogram of the Pdzd8 peptide identified by LC-MS/MS. **D,** Prediction of the interaction between full-length UBA1 and Pdzd8 (CC-domain) by the ZDOCK server. Their structures were visualized using PyMOL software. **E, F,** Co-immunoprecipitation (Co-IP) assays of the interaction between endogenous Pdzd8 and UBA1 in the lysates from NRCMs or mouse hearts, respectively, followed by immunoblotting analysis with anti-Pdzd8 or anti-UBA1 antibodies. Bar: 10 μm. **G,** Fluorescence co-localization of endogenous UBA1 and Pdzd8 proteins in NRCMs. **H,** Identification of the Pdzd8-binding domains within the UBA1. Whole cell lysates isolated from HEK293 cells co-transfected with Flag-tagged full-length and truncated forms of UBA1 and Myc-tagged Pdzd8 plasmids were IP with anti-Myc antibody and analysed by immunoblotting with anti-Flag or Myc antibody. **I,** Schematic diagram for the amino acids (605-843) of the UBA1 binding to Pdzd8. **J,** Mapping of the UBA1-binding domains within the Pdzd8. Whole cell lysates isolated from HEK293 cells co-transfected with Flag-tagged full-length UBA1 and Myc-tagged full-length and truncated forms of Pdzd8 plasmids were IP with anti-Flag antibody followed by immunoblotting with anti-Myc or Flag antibody. **K,** Schematic diagram for the amino acids (1011-1154) of Pdzd8 binding to UBA1. **L,** Lysates from shRNA-NC- or shRNA-UBA1-infected NRCMs following H/R 24 h were IP with anti-Pdzd8 antibody and then analyzed with immunoblotting with anti-ubiquitin (Ub) or anti-Pdzd8 antibody (left). Input for each protein (right). Quantification of ubiquitinated Pdzd8 level (right, n = 3). **M,** Lysates from NRCMs infected with Ad-NC or Ad-Pdzd8 and then following H/R 24 h were IP and then analyzed as described in **L, N,** NRCMs were first infected with adenovirus expressing shRNA-NC or shRNA-UBA1 for 24 h and then stimulated with CHX (20 μM) for another 6, 12, or 24 h. The protein levels of Pdzd8 and UBA1 for each time point were determined by immunoblotting analysis with anti-Pdzd8 or anti-UBA1 antibody (left) and quantification of the ubiquitinated Pdzd8 level (right; n = 3). **O,** NRCMs were infected with Ad-NC or Ad-Pdzd8 and treated and analyzed as described in **n**. Data are expressed as the mean ± SEM, and n indicates the sample number per group. Statistical significance was assessed by two-way ANOVA. *P* values are indicated in the figure. *P* values vs. ShRNA-NC or Ad-NC.

To verify this interaction, structural prediction using ZDOCK suggested UBA1–Pdzd8 binding (Figure 5D). Co-IP in NRCMs using anti-UBA1 or anti-Pdzd8 confirmed reciprocal pull-down of Pdzd8 and UBA1, whereas IgG did not (Figure 5E). Co-IP from WT mouse hearts yielded similar results (Figure 5F). Immunostaining further showed cytoplasmic co-localization of UBA1 and Pdzd8 in NRCMs (Figure 5G). To map the binding sites, we expressed WT and truncated Flag-UBA1 or Myc-Pdzd8 constructs in HEK293 cells. Co-IP using anti-Myc showed that UBA1 aa 605-843 were necessary for binding WT-Pdzd8 (Figure 5H and 5I). Co-IP with anti-Flag showed that Pdzd8 aa 1011-1054 bound WT-UBA1 (Figure 5J and 5K). These findings demonstrate that UBA1 directly interacts with Pdzd8 in cardiomyocytes.

UBA1 initiates ubiquitin activation and transfer to target proteins,^26^ prompting us to examine whether it regulates Pdzd8 protein stability. UBA1^cko^ hearts showed greater I/R-induced Pdzd8 upregulation, whereas UBA1 overexpression reduced this increase (Figure S6B and S6C). We next tested whether UBA1 decreases Pdzd8 by promoting its ubiquitination. IP of Pdzd8 followed by anti-Ub immunoblotting showed that Sh-UBA1 reduced Pdzd8 ubiquitination, whereas Ad-UBA1 enhanced it in NRCMs under control and H/R (Figure 5L and 5M). *In vivo*, Pdzd8 ubiquitination was decreased in UBA1^cko^ hearts and increased in rAAV9-UBA1-injected hearts after I/RI (Figure S6D and S6E), supporting UBA1 as a key mediator of Pdzd8 ubiquitination. We then assessed whether ubiquitinated Pdzd8 undergoes proteasomal degradation. Using CHX pulse-chase, UBA1 knockdown increased Pdzd8 half-life, while UBA1 overexpression reduced it (Figure 5N and 5O). MG132 blocked CHX-induced Pdzd8 loss in Ad-NC and Ad-UBA1 cells (Figure S6F). Together, these findings show that UBA1 promotes Pdzd8 degradation through the ubiquitin-proteasome pathway.

### UBA1 Regulates ER-Mitochondria Contact Sites through Targeting Pdzd8

Because Pdzd8 is a key ER protein required for ERMCS function,^27,28^ we examined whether UBA1 regulates ERMCSs through Pdzd8 *in vitro* and *in vivo*. NRCMs stably expressing Mito-DsRed and Sec61β-EGFP were infected with Ad-GFP, Sh-NC, Ad-UBA1, or Sh-Pdzd8 for 24 h.^29^ Fluorescence imaging showed that ER–mitochondria co-localization increased after H/R in Ad-NC-infected cells but was reduced by Ad-UBA1 (Figure 5A). Conversely, Sh-UBA1 further enhanced H/R-induced ER–mitochondria co-localization compared with Sh-NC, and this increase was attenuated when Sh-Pdzd8 was present (Figure 6B), indicating that UBA1 suppresses Pdzd8-dependent ERMCS formation.

**Figure 6.**
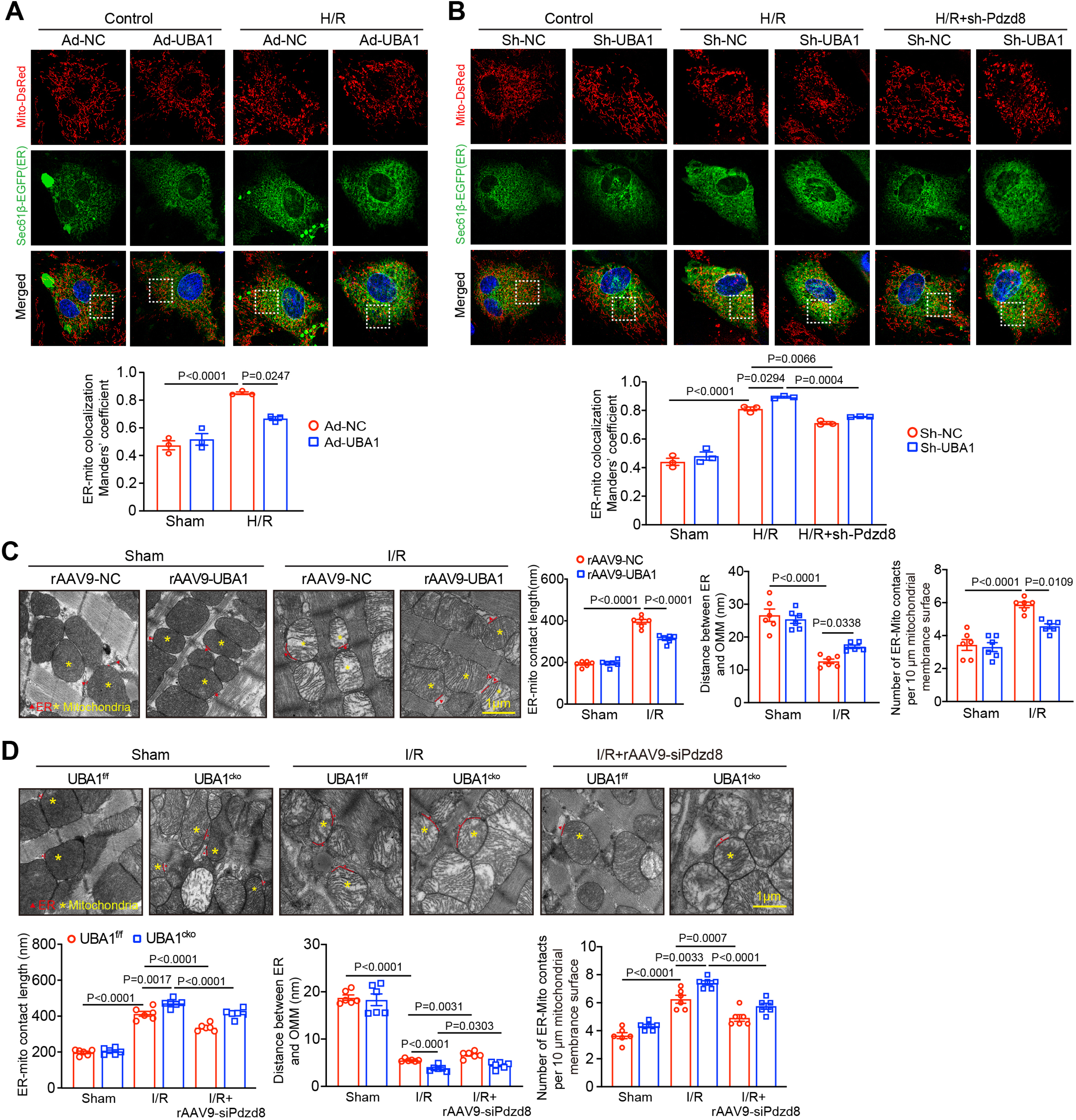
UBA1 reduces ER-mitochondria contacts through regulating Pdzd8. **A,** NRCMs were infected with Sec61β-EGFP (an ER marker, green) and/or Mito-DsRed (a mitochondrial marker, red) for 24 h and then infected with adenovirus expressing UBA1 (Ad-UBA1) or Ad-NC control for an additional 24 h in the presence or absence of H/R. Representative images of the co-localization of ER and mitochondria captured by a structured illumination microscope (SIM) (top). Bar: 10 μm. Quantitative analysis of ER-mitochondria contacts using the Mander coefficient by ImageJ. n = 15-20 mitochondria examined over 3 independent experiments. **B,** NRCMs were infected with Sec61β-EGFP and/or Mito-DsRed for 24 h and then infected with adenovirus expressing sh-NC, sh-UBA1, together with sh-Pdzd8 in the presence or absence of H/R. Representative image by analyzed structured illumination microscope (SIM) and quantification of ER-mitochondria contacts as described in **A**. **C,** Representative TEM images of ER-mitochondria contacts in Sh-NC, Sh-UBA1, or Sh-Pdzd8 infected NRCMs, with mitochondria in red triangle, ER in yellow asterisk, and MAMs in red dotted line (top, right). The mitochondrial length, the distance between ER and OMM, and the number of ER-Mito contacts per 10 μm mitochondrial membrane surface were further analyzed by ImageJ software (n = 6, bottom, right). Bar: 1 cm. **D,** Representative TEM images of ER-mitochondria contacts in Ad-NC or Ad-UBA1 infected NRCMs, with mitochondria in red triangle, ER in yellow asterisk, and MAMs in red dotted line (left). Bar: 1 cm. The analyzed as described in “c” (n = 6, right). Data are expressed as the mean ± SEM, and n indicates the independent experiments or animal samples per group. Data are expressed as the mean ± SEM, and n indicates the sample number per group. Statistical significance was assessed by two-way ANOVA. *P* values are indicated in the figure.

*In vivo*, WT mice received rAAV9-NC or rAAV9-UBA1, and UBA1^f/f^ or UBA1^cko^ mice received rAAV9-si-Pdzd8 for 3 weeks, followed by I/RI for 24 h. TEM showed that I/R increased ERMCS formation, reduced ER–outer mitochondrial membrane distance, increased contact length, and more ER–mito contacts in rAAV9-NC or UBA1^f/f^ hearts, while rAAV9-UBA1 reduced these changes (Figure 6C). I/R-mediated ERMCS formation was further enhanced in UBA1^cko^ hearts but abolished in UBA1^f/f^ and UBA1^cko^ mice treated with rAAV9-siPdzd8 (Figure 6D), identifying Pdzd8 as the key ERMCs regulator. These findings show that UBA1inhibits I/R-induced ER–mitochondria interactions by reducing Pdzd8 stability.

### Cardiomyocyte-Specific Pdzd8 Knockout Alleviates Myocardial Injury and Dysfunction *in vivo and in vitro*

To evaluate the role of Pdzd8 in cardiomyocytes, WT mice were injected with rAAV9-siPdzd8 or rAAV9-siNC for 3 weeks and then subjected to sham or I/R for 24 h (Figure S7A). Pdzd8 knockdown reduced endogenous Pdzd8 protein by 40-70% without altering UBA1 (Figure S7B). Echocardiography showed that Pdzd8 silencing improved I/R-induced cardiac dysfunction, increasing EF% and FS% (Figure S7C, Table S6). rAAV9-siPdzd8 also reduced infarct size, TUNEL+ myocytes, and ROS generation compared with rAAV9-siNC (Figure S7D through 7F). Correspondingly, I/R-induced increases in NOX2/NOX4 mRNAs, serum LDH, and Bax protein, and decreases in Bcl-2 were reversed by Pdzd8 knockdown (Figure S7G through 7I), indicating cardioprotection *in vivo*.

Given Pdzd8’s role in ERMCSs (Figure 6), Ca^2+^ handling, mitochondrial function, and ER stress, we examined its loss in NRCMs exposed to H/R. After 24 h, Sh-Pdzd8 (MOI: 50) suppressed H/R-induced apoptosis, ROS, Ca^2+^ release, mitochondrial fragmentation, and loss of MMP compared with Sh-NC (Figure S8A through 8E). Sh-Pdzd8 also decreased Drp1 and increased Mfn-1/2 post-H/R (Figure S8F). Together, these results show that Pdzd8 knockdown protects cardiomyocytes against I/R-related injury and dysfunction.

### Cardiac-Specific Deletion of UBA1 Aggravates Myocardial I/RI by increasing Pdzd8 Stability

Because UBA1 interacts with Pdzd8 and UBA1 loss reduces Pdzd8 ubiquitination and degradation (Figure 5), increasing ERMCS and promoting mitochondrial dysfunction and ER stress (Figure 6 and 7), while Pdzd8 knockdown alleviates myocardial I/RI *in vivo* and *in vitro* (Figure S7 and S8), we hypothesized that UBA1 deficiency aggravates I/RI through Pdzd8. To test this, UBA1^f/f^ and UBA1^cko^ mice received rAAV9-siPdzd8 or rAAV9-siNC for 3 weeks before I/R for 24 h. rAAV9-siPdzd8 reduced Pdzd8 by nearly 80% without affecting UBA1 (Figure S9). As in Figure 2, UBA1^cko^ mice treated with rAAV9-siNC displayed worse cardiac dysfunction, larger infarcts, more TUNEL^+^ myocytes, higher ROS, and an increased Bax/Bcl-2 ratio compared with UBA1^f/f^ (Figure 7A through 7E; lane 2 vs. 1, Table S7). These effects were markedly alleviated in UBA1^cko^ mice receiving rAAV9-siPdzd8 (lane 4 vs. 2), with similar protection also observed in UBA1^f/f^ mice (lane 3 vs. 1). Pdzd8 knockdown also reduced Drp1, increased Mfn-1/2, and enhanced ETC complex I/II activities and ATP levels compared with rAAV9-siNC in both genotypes after I/R (Figure 7F and 7G). ER stress markers p-IRE1α, p-PERK, and ATF6 were also lower with rAAV9-siPdzd8 (Figure 7H). These findings show that the deleterious phenotype of UBA1 deficiency in I/R injury is mediated by elevated Pdzd8 *in vivo*.

**Figure 7.**
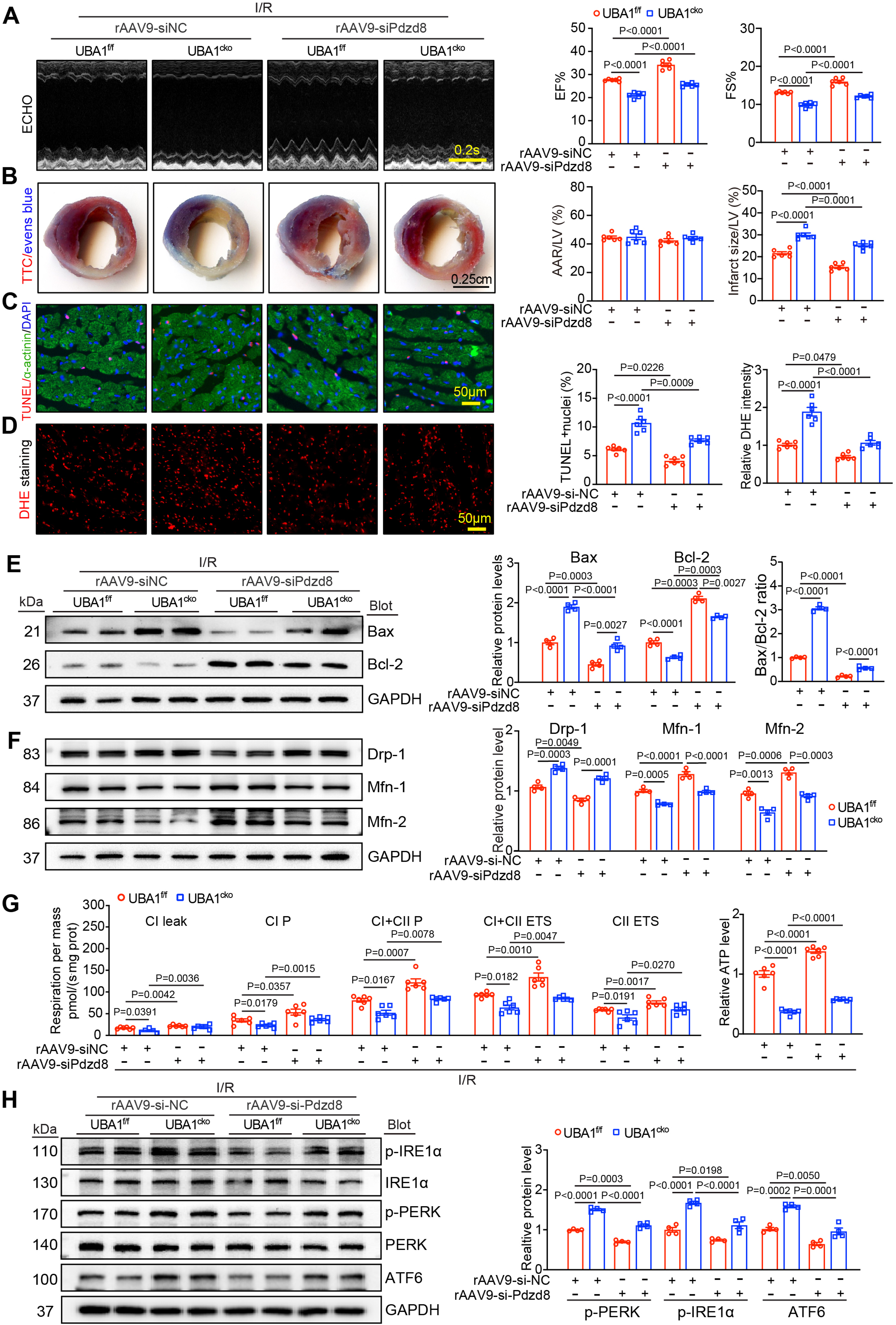
Knockdown of Pdzd8 in UBA1^cko^ mice reverses I/R-induced cardiac damage and dysfunction. **A,** Male UBA1^f/f^ or UBA1^cko^ mice were injected with rAAV9-siNC or rAAV9-siPdzd8 (1.0×10^12^ vg/mg) for 3 weeks and then exposed to I/RI for another 24 h. Representative echocardiographic images of the LV function (left), and the analysis of the LV EF% and LV FS% (right, n = 6). Bar: 0.2 s. **B,** TTC–Evans blue staining of cardiac infarct areas, and the ratios of the AAR to the LV area and the infarct area relative to the LV area (n = 6). Bar: 0.5 cm. **C,** Representative images of TUNEL staining (red) in the border-zone of cardiac infarct areas (left). Immunostaining of myocardial myocytes and nuclei with anti-α-actinin (green) and DAPI (blue), respectively (left). The percentage of TUNEL-positive nuclei (right, n = 6). Bar: 50 μm. **D,** Immunostaining of heart sections with DHE dye (left), and quantification of ROS levels (right, n = 6). Bar: 50 μm. **E, F,** Immunoblotting analysis of Bax, Bcl-2, Drp1, Mfn1, and Mfn2 protein levels (left), and quantification of each protein intensity (right, n = 4). **G,** Summary data from O2K measurements of respiratory capacity, including CI leak, CI P, CI + CII P, CI + CII ETS, CII ETS, and ATP production (n = 6). **H,** Immunoblotting analysis of p-IREɑ, IREɑ, p-PERK, PERK, and ATF6 protein levels (left), and quantification of them (right, n = 4). Data are expressed as the mean ± SEM, and n indicates the sample number per group. Statistical significance was assessed by two-way ANOVA. *P* values are indicated in the figure.

### Auranofin induces UBA1 activity to Prevent Myocardial Injury in I/R Mouse and H/R cardiomyocyte Models

Because UBA1 overexpression improved myocardial function (Figure 4), we hypothesized that activating UBA1 could protect against myocardial I/RI. Auranofin (AF), a gold compound used clinically for rheumatoid arthritis, was recently identified as a UBA1 activator (Figure 8A).^30,31^ To evaluate whether AF protects against I/RI, WT mice were treated with AF (5-40 mg/kg) for 24 h (Figure 8B). LDH levels showed that 20 mg/kg AF was non-cytotoxic (Figure 8C) and reduced Pdzd8 expression without altering UBA1 protein (Figure 8D), identifying 20 mg/kg as an appropriate dose. WT mice then received AF 24 h before I/R. AF improved cardiac dysfunction, reduced infarct size, decreased TUNEL^+^ cells, and lowered the Bax/Bcl-2 ratio compared with vehicle (Figure 8E through 8H, Table S8). In contrast, AF did not rescue cardiac dysfunction or infarct size in AF-treated UBA1^cko^ mice (Figure S10A and S10b, Table S9), demonstrating that AF requires UBA1 to exert protection. AF also corrected I/R-induced mitochondrial abnormalities and excessive ER stress, reducing Drp1, p-IRE1α, p-extracellular signal-regulated kinase, and ATF6 while restoring Mfn-1/2 in WT hearts (Figure 8I). *In vitro*, AF reduced H/R-induced ER–mitochondria colocalization in NRCMs (Figure 8J). These findings indicate that AF protects against myocardial I/RI by activating UBA1 and reducing Pdzd8-dependent ERMCS formation.

**Figure 8.**
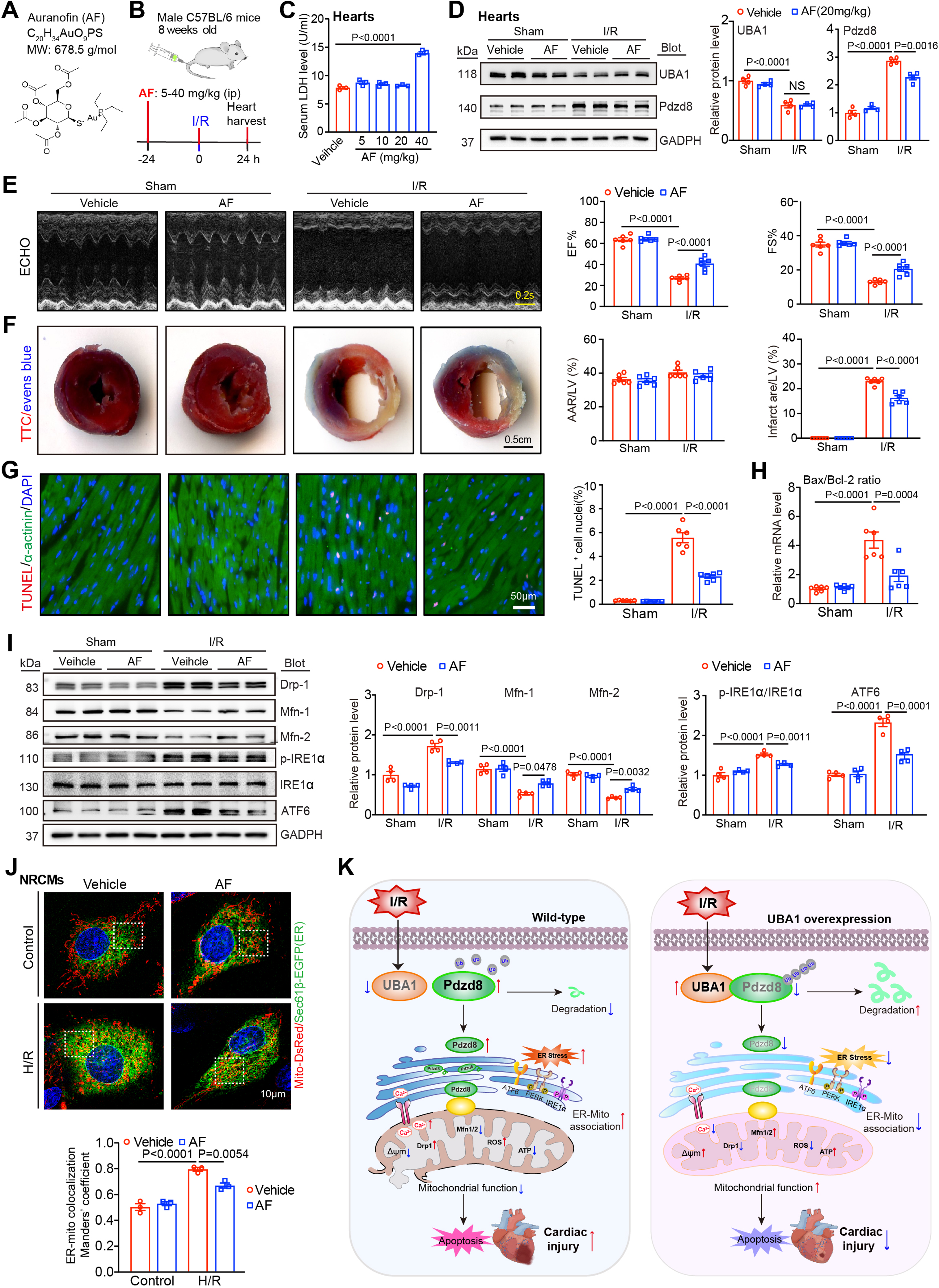
Auranofin enhances UBA1 activity to alleviate myocardial I/RI in mice. **A,** Chemical structure of Auranofin (AF). **B,** Schematic diagram of the experimental design. **C,** Measurement of serum LDH level in mice (n = 3) treated with different dosages (0, 10, 20, or 40 mg/kg) of AF. **D,** Immunoblotting analysis of UBA1 and Pdzd8 protein levels in heart infarct tissue of mice treated with 20 mg/kg of AF post-I/R (left), and quantification of relative protein levels (right, n = 4). **E,** Representative echocardiographic images of the LV function (left), and the analysis of the LV EF% and LV FS% (right, n = 6). Bar: 0.2 s. **F,** Representative images of TTC-Evens blue staining of the infarct area post-I/R (left). Ratios of AAR to LV area and infarct size to LV area (right, n = 6). Bar: 2.5 cm. **G,** Representative TUNEL staining images of the infarct border zone (left), and the percentage of TUNEL-positive nuclei (right, n = 6). Bar: 50 μm. **H,** qPCR analysis of Bax and Bcl-2 mRNA levels in the infarct border zone (n = 6). **I,** Immunoblotting analysis of Drp1, Mfn-1, Mfn-2, p-IRE-ɑ, IREɑ, and ATF6 protein levels in the infarct border zone (left), and quantification of them (n = 4). **J,** NRCMs were infected with Sec61β-EGFP (green) and Mito-DsRed (red) for 24 h and then treated with vehicle or AF (200 nm) for another 24 h in the presence or absence of H/R. Representative SIM images of the co-localization of ER and mitochondria (left). Bar: 10 μm. Quantitative analysis of ER-mitochondria contacts using the Mander coefficient by ImageJ (n = 15-20 mitochondria examined over 3 independent experiments). **K.** Mechanism diagram illustrating the crucial role of the UBA1-Pdzd8 pathway in regulating ER-mitochondria contacts and myocardial I/R. Data are expressed as the mean ± SEM, and n indicates the sample number per group. Statistical significance was assessed by two-way ANOVA. *P* values are indicated in the figure.

To further confirm UBA1 activation as protective *in vitro*, AF at 200 nM showed no cytotoxicity and did not alter UBA1 expression, yet markedly increased UBA1 activity in NRCMs, as evidenced by enhanced total protein ubiquitination (Figure S11A through S11C). AF treatment for 24 h significantly reduced H/R-induced TUNEL-positive cells, Ca^2+^ release, mitochondrial fission, and ATP depletion compared with vehicle (Figure S11D through S11G). These benefits were completely lost in Sh-UBA1-infected NRCMs, indicating the requirement of UBA1 for AF’s protective effects. Collectively, these findings show that AF prevents H/R-induced cardiomyocyte injury by activating UBA1, supporting its potential as a therapeutic option for myocardial I/RI.

## DISCUSSION

In this study, we identify UBA1 as a key regulator of myocardial I/RI through its control of ER–mitochondria coupling via Pdzd8 ubiquitination. UBA1 expression was reduced in cardiomyocytes of I/R mice and MI patients and manipulating UBA1 levels either exacerbated or mitigated I(H)/R-induced injury *in vivo* and *in vitro*. Interactome analysis revealed Pdzd8 as a major UBA1 substrate. Mechanistically, UBA1 binds Pdzd8 and promotes its ubiquitination and degradation, thereby limiting ERMCS formation and reducing mitochondrial dysfunction, ER stress, and Ca^2+^ transfer during reperfusion. Systemic activation of UBA1 also protected against myocardial I/RI. Overall, these findings establish UBA1 as a central regulator of ERMCS dynamics through Pdzd8 degradation and identify a previously unrecognized therapeutic mechanism for myocardial I/RI (Figure 8K).

Protein homeostasis is essential for cardiac function. Ubiquitination, a major PTM, regulates protein degradation and contributes to numerous cardiac disorders. Many components of the ubiquitin-proteasome system (UPS), including E3 ligases, DUBs, and proteasome catalytic subunits, influence immune responses, myocardial I/RI and MI by modifying distinct substrates.^32–36^ Our recent observations showed that Psmb8 and Psmb10 protect the heart during I/RI by modulating mitochondrial dynamics through Drp1 and Parkin/Mfn1/2 pathways,^37,38^ suggesting that UPS enzymes may represent therapeutic targets for ischemic heart disease. However, the role of E1 enzymes in myocardial I/RI is unclear. Only two E1s, UBA1 and UBA6, activate ubiquitin in humans. UBA1 is implicated in VEXAS syndrome, cancer, neurodegeneration, atherosclerosis, restenosis, and cardiac remodeling,^11–14^ yet its involvement in myocardial I/RI has been undefined. Here, we show that UBA1 expression is markedly reduced in mouse I/R and human MI hearts, especially in H/R-stimulated cardiomyocytes (Figure 1). Loss of UBA1 caused severe myocardial injury, mitochondrial dysfunction, ER stress, and abnormal Ca^2+^ handling, all of which were worsened in UBA1^cko^ mice (Figure 2 and 3) and alleviated by UBA1 overexpression (Figure 4). These findings identify cardiomyocyte-derived UBA1 as a key regulator of myocardial I/RI through its influence on ER–mitochondria homeostasis.

Extensive evidence shows that mitochondrial dysfunction, ER stress, ROS generation, and impaired calcium homeostasis are major drivers of myocardial I/RI pathology.^39,40^ ERMCSs form a dynamic interface between the ER and mitochondria that governs Ca^2+^ transfer, lipid exchange, ER stress, and mitochondrial dynamics.^39,40^ More than 1000 MAM-associated proteins, including IP3R, GRP75, VDAC1, PTPIP51, Drp1, Mfn2, and Pdzd8, maintain ERMCS integrity, and their dysfunction contributes to diverse diseases.^41^ Restoring ERMCS balance can reduce mitochondrial dysfunction and ER stress, limiting cardiomyocyte death and attenuating myocardial I/RI and MI progression.^42^ For example, PERK activation by ZY341 disrupts VAPB–PTPIP51 binding, reducing ERMCS-mediated Ca^2+^ and phosphatidic acid transfer and improving mitochondrial dynamics after I/R.^43^ In addition, cardiac GSK-3β localizes to the sarcoplasmic reticulum (SR)/ER and ERMCSs, and its inhibition reduces I/R-induced cytosolic and mitochondrial Ca^2+^ overload and cell death by interacting with IP3R.^44^ Several ERMCS-associated proteins influence myocardial I/RI. Mtus1A supports ERMCS integrity by enhancing IP3R1-Grp75-VDAC assembly and protects against myocardial injury.^20^ TBC1D15 interacts with Drp1 at mitochondria–lysosome contact sites to limit mitochondrial division after I/R, while DIAPH1–MFN2 coupling decreases ER–mitochondria distance and reduces I/RI susceptibility.^45^ Conversely, PTPIP51 inhibition protects the heart by reducing SR–mitochondria contacts and limiting mitochondrial Ca^2+^ uptake.^46^ Among these proteins, Pdzd8 is an ER-resident tether essential for ERMCS stability via FKBP8 interaction.^20^ Loss of Pdzd8 disrupts Ca^2+^ signaling, lipid transport, autophagy, apoptosis, and mitochondrial and endosomal function, contributing to neurological, metabolic, and renal disorders in mice.^24,25,27,28,47–51^ Its role in myocardial I/RI, however, was unknown. We show that Pdzd8 expression increases in I/R mouse hearts and H/R-stimulated NRCMs (Figure S7 and S8), whereas Pdzd8 knockdown improves cardiac function, reduces infarction, ROS, Ca^2+^ release, ER stress, mitochondrial dysfunction, and ERMCS formation in I/R or H/R conditions (Figure S7 and S8). These findings identify Pdzd8 as a pathological driver of myocardial I/RI and reveal new insights into its role in reperfusion injury.

ER–mitochondria contact stability is essential for Ca^2+^ and lipid transfer. Pdzd8 is a critical ERMCS tether, but mechanisms regulating its stability remain unclear. AMPK was recently shown to activate Pdzd8 and glutaminase 1 (GLS1) through Pdzd8 T527 phosphorylation to promote glutaminolysis.^52^ Our in vivo and in vitro findings show that Pdzd8 protein increases in cardiomyocytes after I/R, but how ubiquitination controls its stability at ERMCSs has been unclear. Here, we identify UBA1-mediated ubiquitination as a PMT mechanism regulating Pdzd8 stability during myocardial I/RI. Interactome analysis, ZDOCK prediction, Co-IP/MS, and immunostaining reveal Pdzd8 as a new UBA1-binding protein in cardiomyocytes, with interaction between the UBA1 UAE domain (605-843 aa) and Pdzd8 CC-domain (1011-1154 aa) (Figure 5). Previous work shows UBA1 modifies proteins such as TRAF6, IĸB-α, NF-ĸB, GRP78, and ATF4.^11–14^ We now identify Pdzd8 as an additional target. K48-linked ubiquitin chains direct proteins to the 26S proteasome, whereas K63-linked chains regulate DNA repair and signaling.^53^ Zebrafish UBA1 degrades interferon (IFN) regulatory factor 3 (IRF3) through K48-linked ubiquitination to limit IFN production.^54^ Consistent with this, we show UBA1 drives Pdzd8 degradation via K48-linked polyubiquitination, decreasing Pdzd8 accumulation and ERMCS formation (Figure 5, Figure S6). Moreover, Pdzd8 knockdown markedly reduces ERMCS density, Ca^2+^ and ROS release, mitochondrial dysfunction, ER stress, and myocardial I/RI in UBA1^cko^ mice (Figure 6 and 7). Together, these results establish UBA1-directed K48-linked ubiquitination of Pdzd8 as a key mechanism in myocardial I/RI.

Growing evidence shows that multiple UPS enzymes are viable therapeutic targets, and several UPS activators or inhibitors improve ischemic heart disease.^35,55^ We recently identified proteasome activators such as ursolic acid, MK-886, TCH-165, and AM404 that mitigate myocardial I/RI by increasing immunoproteasome subunit expression and activity,^56–59^ underscoring the therapeutic relevance of UPS modulation in I/RI and post-MI heart failure. Auranofin, an FDA-approved gold compound used for rheumatoid arthritis, was recently shown to activate UBA1 and promote ubiquitination and clearance of misfolded ER proteins.^31^ Beyond its anti-inflammatory and antioxidant actions, auranofin is being evaluated for COVID-19, cancer, neurodegeneration, and hepatic fibrosis.^60–63^ Although high-dose auranofin (100 mg/kg) increases cell apoptosis and I/RI in isolated rat hearts,^64^ our findings show that low-dose auranofin (20 mg/kg) markedly reduces I/R-induced ERMCS formation, mitochondrial dysfunction, ER stress, and cardiomyocyte apoptosis by enhancing UBA1 activity, thereby improving cardiac injury and function (Figure 8, Figure S11). These results extend the therapeutic potential of auranofin to ischemic heart disease.

This study has some limitations. We did not determine how I/R lowers UBA1 expression in cardiomyocytes, identify the E3 ligases responsible for Pdzd8 K48-linked ubiquitination, or define the tethering partners that cooperate with Pdzd8 to regulate ERMCS structure and function. The long-term role of UBA1 in post-MI remodeling also requires further investigation. Although activating UBA1 shows therapeutic promise for myocardial I/RI, its applicability in human ischemic disease remains uncertain.

In conclusion, this study shows that UBA1 protects cardiomyocytes during I/RI by preserving ERMCS structure and function through Pdzd8 ubiquitination. Increased UBA1 enhances Pdzd8 K48-linked polyubiquitination and proteasomal degradation, reducing ERMCS formation and subsequently limiting mitochondrial dysfunction and ER stress, thereby alleviating myocardial I/RI. These findings identify UBA1-Pdzd8 signaling as a previously unrecognized contributor to myocardial I/RI, and suggest that enhancing UBA1 activity or reducing Pdzd8 may offer a potential therapeutic strategy.

## Nonstandard Abbreviations and Acronyms

AAV9: adeno-associated virus serotype 9
AF: Auranofin
CKO: cardiomyocyte-specific UBA1 knockout
Co-IP: Co-immunoprecipitation
DEGs: differentially expressed genes
DUBs: deubiquitinases
ECHO: echocardiography
ERMCs: endoplasmic reticulum–mitochondria contacts
ERMCSs: endoplasmic reticulum-mitochondrial contact sites
EF: ejection fraction
FS: fractional shortening
H/R: hypoxia/reoxygenation
I/RI: ischemia reperfusion injury
LDH: lactate dehydrogenase
MAMs: mitochondria-associated ER membranes
MI: myocardial infarction
NRCFs: neonatal rat fibroblasts
NRCMs: neonatal rat cardiomyocytes
Pdzd8: PDZ domain containing 8
ROS: reactive oxygen species
TEM: transmission electron microscopy
TUNEL: TdT-mediated dUTP nick-end labeling
Ub: Ubiquitin
UBA1: Ubiquitin-activating enzyme E1
WT: wild type

## Author Contributions

L-L. Xu and P-B. Li performed the experiments and analyzed the data. P-B. Li performed animal surgery, infarct staining, and echocardiography. L-L. Xu performed protein interaction. W-X. Jiang performed bioinformatics analysis. H-H. Li and L-L. Xu conceived and designed the research. H-H. Li, W-X. Jiang and J. Du supervised the research. H-H. Li, J. Du and W-X. Jiang drafted the manuscript. H-H. Li provided funding. All authors reviewed and approved the manuscript.

## Sources of Funding

This work was supported by a grant from the National Natural Science Foundation of China (No. 82030009).

## Disclosures

None

## Supplemental Material

Expanded Materials and Methods

Figures S1–S11

Tables S1–S9

References 65–67

## Notes

### Competing Interest Statement

The authors have declared no competing interest.

